# In vitro blood clot mechanical properties depend on fibrinogen and white blood cell subtypes in addition to hematocrit

**DOI:** 10.1101/2025.10.28.685168

**Authors:** Grace N. Bechtel, Nandini Senthilkumar, Juliette Noyer, Tatum Eades, Jan Fuhg, Jonathan I. Tamir, Hamidreza Saber, Sapun H. Parekh, Manuel K. Rausch

**Affiliations:** Department of Biomedical Engineering, The University of Texas at Austin, Austin, TX, United States of America; Department of Aerospace Engineering and Engineering Mechanics, The University of Texas at Austin, Austin, TX, United States of America; The Oden Institute for Computational Engineering and Sciences, The University of Texas at Austin, Austin, TX, United States of America; Department of Electrical and Computer Engineering, The University of Texas at Austin, Austin, TX, United States of America; Department of Diagnostic Medicine, Dell Medical School, The University of Texas at Austin, Austin, TX, United States of America; Department of Neurology and Neurosurgery, Dell Medical School, The University of Texas at Austin, Austin, TX, United States of America; Department of Mechanical Engineering, The University of Texas at Austin, Austin, TX, United States of America

**Author notes:** Corresponding Author: Manuel Rausch, 2617 Wichita Street, Austin, TX 78712.

**Keywords:** Thrombus, Hematocrit, Fibrinogen, Stiffness, Fracture Toughness

## Abstract

**Background:** Understanding the role of blood composition in clot mechanics may provide critical cues toward their diagnosis and treatment. To this end, we previously showed that sex and standard blood composition measures explain some variability in clot mechanics, but much remains unaccounted for. Ours and others’ studies ignored the roles of fibrinogen and white blood cell (WBC) subtypes, which is surprising given their physiological importance.

**Objective:** To develop a more complete understanding of what determines clots’ mechanical properties, we now study the role of previously untested factors, namely fibrinogen levels and WBC subtypes, in addition to sex and standard blood composition measures.

**Methods:** We drew blood from healthy young adults and prepared in vitro clot samples. Using pure shear and mode-I fracture tests, we measured clots’ stiffness, fracture toughness, strength, and work to rupture. We then used linear regressions to quantify how blood composition influences clot mechanical properties, reporting R^2^ to assess explanatory power.

**Results:** We found that fibrinogen is the strongest predictor of clot mechanical properties, positively correlating with all four metrics and achieving the highest average R^2^. For example, fibrinogen accounts for 70% of the variation in stiffness and 78% in fracture toughness. Among WBC subtypes, neutrophils positively correlate with strength and work to rupture, whereas eosinophils negatively correlate with strength.

**Conclusion:** Our work shows that fibrinogen levels and WBC subtypes are previously untested but major determinants of in vitro clot mechanical properties. Future studies should connect these findings to in vivo clot behavior and clinical outcomes.

## 1. Introduction

The mechanical properties of blood clots play a key role in their physiological function and influence the development and severity of thromboembolic disease [1]. For example, clot mechanical properties impact embolization risk, determine where a clot may lodge in the vasculature, and influence how a clot responds to treatments such as thrombolysis and mechanical thrombectomy [2, 3]. Thus, understanding clot mechanical properties is critical for guiding thromboembolic disease prevention and treatment.

Since clots cannot be directly accessed or tested in vivo, we and others commonly use in vitro clots as models [4–6]. These models enable controlled measurements of clot properties that are not possible in vivo. Importantly, prior studies suggest that the structure of a patient’s in vitro clot reflects their in vivo clot behavior and thrombotic risk [7–9]. Based on these findings, we expect that in vitro clot mechanical properties are similarly informative for understanding clot behavior and thrombotic risk in vivo.

In our prior work, we tested the sensitivity of in vitro blood clot mechanical properties to sex, age, and complete blood count (CBC) data in a study population spanning ages 19-46 [10]. We found that sex had a strong effect on in vitro clot mechanical properties, blood composition measures had a moderate impact, and age had no significant impact. These findings raised important questions. For instance, hematocrit, widely considered a primary driver of clot mechanics [4, 5], did not account for a large amount of the observed variation in mechanical properties. This highlights the presence of additional contributing factors not included in our study. Furthermore, our analysis suggested that white blood cell (WBC) count influences clot mechanical properties, but without differential counts, the mechanisms by which specific WBC subtypes affect clot properties remain unclear.

Thus, our goal is to build on our prior work to identify what blood components drive variability in the mechanical properties of in vitro clots. To this end, we reduced the age range of our study population to healthy adults in their 20s, mitigating potential sources of variability unrelated to blood composition. We also expanded our blood composition analysis to include fibrinogen levels and white blood cell subtypes in addition to basic CBC data. We evaluated clot stiffness, fracture toughness, strength, and work to rupture to capture their elastic and fracture properties. We then correlated these measures with subjects’ blood composition data to identify the key drivers of clot mechanical properties.

## 2. Methods

### 2.1. Blood Collection

We collected venous blood samples from six healthy female subjects (ages 22-25) and six healthy male subjects (ages 20-27) under Institutional Review Board approval at The University of Texas at Austin. All subjects provided written informed consent. We drew blood into acid citrate dextrose (ACD) anticoagulant and prepared and tested clot samples within three hours of collection [11]. We also collected blood into dipotassium ethylenediaminetetraacetic acid (*K*_2_*EDTA*) and trisodium citrate tubes. We submitted these samples to a local clinical laboratory for CBC with differential and a fibrinogen assay. These tests provide standard measures of red blood cells, detailed white blood cell counts, and fibrinogen levels. We report subject sex, age, and blood data in Supplementary Table 1.

### 2.2. Sample Preparation

We created whole blood clots using our previously established protocol [12, 13]. To reverse the effects of the anticoagulant and initiate coagulation, we added 10% (w/v) calcium chloride to blood samples to a final concentration of 20 mM. We gently mixed the solution and cast it into Velcro-lined, 3D-printed molds (40 × 10 × 3 mm). We covered the samples and incubated them for 60 minutes at 37°C. From each volunteer, we prepared six samples, providing three technical replicates per test modality.

### 2.3. Mechanical Testing

We performed extension-to-failure tests in pure shear and mode-I fracture geometries [14, 15]. To prepare mode-I samples, we cut a 13 mm lateral notch. Then, we mounted samples on our uniaxial tensile tester (Instron, Norwood, MA, USA) and extended them until failure at a strain rate of 0.2 mm/s. Throughout the test, we measured force with a 10 N load cell (accuracy of ± 0.02 N) and logged displacement data. We also captured images of the samples at a rate of 5 Hz.

Using data from pure shear tests, we calculated clot stiffness, strength, and work to rupture. To quantify stiffness, we calculated the tangent modulus of each sample at a stretch ratio of 1.35. A stretch ratio of 1.35 was chosen to provide a consistent reference across samples and capture the clots’ material response prior to damage [4, 10]. We measured strength as the peak stress of each sample and work to rupture as the total energy absorbed before failure. To calculate fracture toughness, we used images of the sample during mode-I tests to identify *λ*_*c*_, the stretch at which crack propagation begins. We then computed fracture toughness as

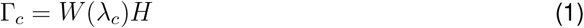

where H is the initial sample height (10 mm) and *W*(*λ*_*c*_) is the strain energy density (i.e., the area under the stress-stretch curve) of the corresponding pure shear sample [16, 17].

### 2.4. Statistics

We performed all statistical analyses in R (The R Foundation, Vienna, Austria, Version 4.3.2). First, we used independent, two-sided Student’s t-tests to evaluate sex differences in blood composition metrics and clot mechanical properties. To account for multiple comparisons, we applied Holm-Bonferroni corrections separately to each group, and all reported p-values reflect these adjustments. We present results as mean ± standard deviation and define statistical significance as adjusted p *<* 0.05.

Next, we examined relationships between clot mechanical properties and CBC measures. For each subject, we averaged mechanical property values across three technical replicates to obtain a single measurement per metric. We then calculated Pearson’s correlation coefficients and visualized the results as a heatmap. Given the large number of correlations tested, we applied the Benjamini-Hochberg procedure to control the false discovery rate. All reported p-values reflect these corrections. For CBC measures that showed more than one significant correlation, we plotted linear regression fits with 95% confidence intervals.

## 3. Results

### 3.1. Blood composition differs by sex

Figure 1 summarizes the blood composition metrics of our subject population split by sex. Our results are consistent with known sex-related trends in red blood cell metrics. For example, male subjects exhibit higher hematocrit, RBC count, and hemoglobin compared to female subjects, indicating greater oxygen-carrying capacity (47.97 ± 2.74 vs. 40.03 ± 3.75 %, p = 0.019; 5.32 ± 0.15 vs. 4.38 ± 0.37 M/uL, p = 0.002; and 16.02 ± 0.83 vs. 13.18 ± 1.42 g/dL, p = 0.019, respectively), see Figure 1A-C.

**Figure 1:**
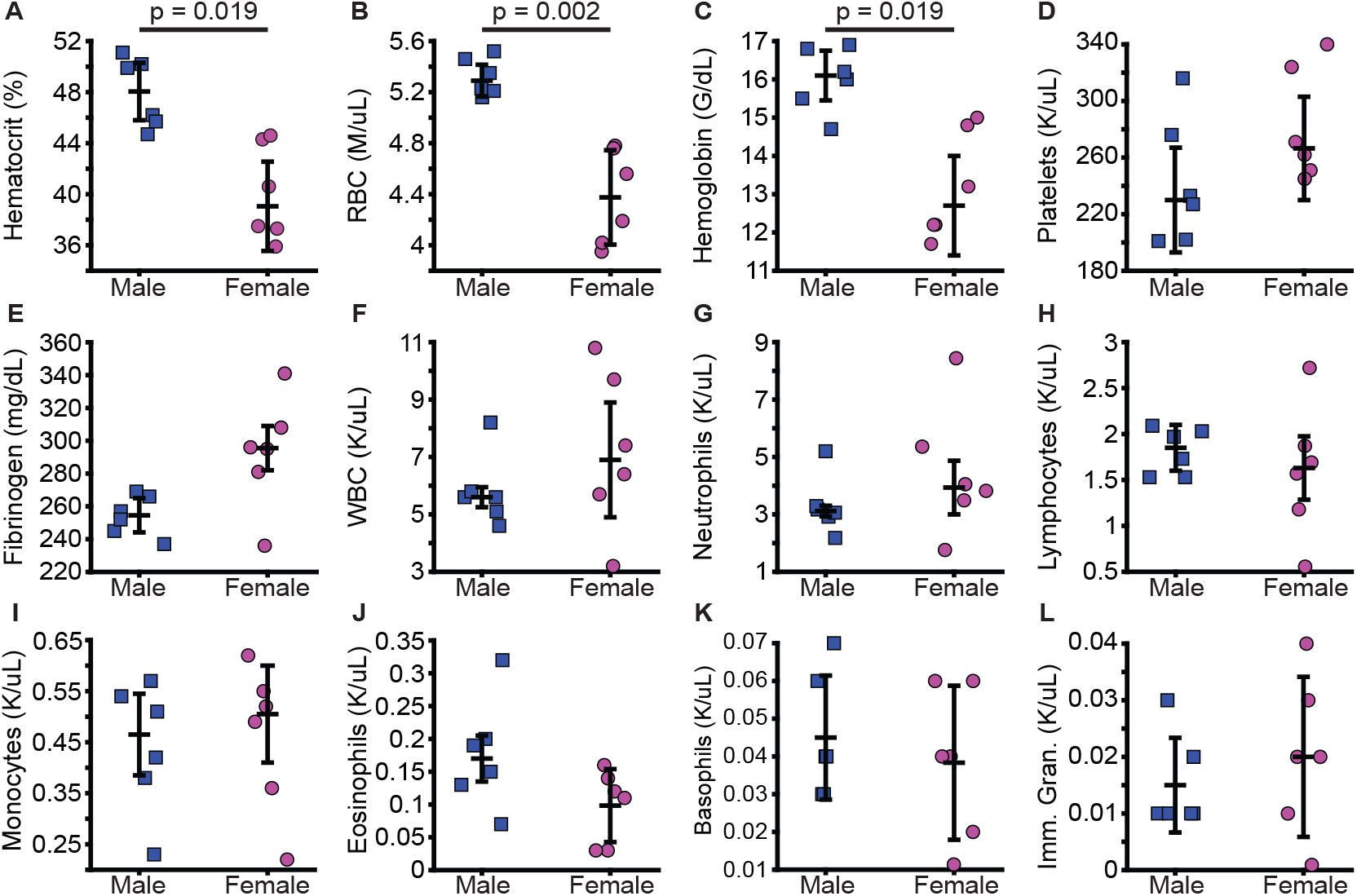
Blood composition by sex. (A-C) Male subjects have higher hematocrit, RBC counts, and hemoglobin compared to female subjects. (D-L) There are no significant sex differences in platelet count, fibrinogen levels, total WBC count, or any WBC subtype, including neutrophils, lymphocytes, monocytes, eosinophils, basophils, or immature granulocytes. Each point represents data from one subject, and error bars indicate mean ± 1 standard deviation.

We did not observe sex differences in other blood composition metrics. Platelet counts are similar between male and female subjects (242.50 ± 45.21 vs. 282.17 ± 39.95 K/uL, p = 0.966), see Figure 1D. Male and female subjects do not have significantly different fibrinogen levels (254.33 *±* 12.26 vs. 292.83 *±* 34.45 mg/dL, p = 0.243), see Figure 1E. Similarly, male and female subjects have comparable total WBC counts (5.82 *±* 1.25 vs. 7.20 ± 2.76 K/uL, p > 0.999) and WBC subtype counts, including neutrophils (3.30 ± 1.10 vs. 4.49 ± 2.26 K/uL, p > 0.999), lymphocytes (1.81 ± 0.25 vs. 1.59 ± 0.74 K/uL, p > 0.999), monocytes (0.44 ± 0.13 vs. 0.46 ± 0.15 K/uL, p > 0.999), eosinophils (0.18 ± 0.08 vs. 0.10 ± 0.06 K/uL, p = 0.696), basophils (0.05 ± 0.02 vs. 0.04 ± 0.02 K/uL, p > 0.999), and immature granulocytes (0.02 ± 0.01 vs. 0.02 ± 0.01 K/uL, p > 0.999), see Figure 1F-L.

### 3.2. Blood clot mechanical properties differ between male and female subjects

Figure 2A-B show the Cauchy stress-stretch curves under pure shear and mode-I loading, and Figure 2C-F show the resulting scalar metrics. We first assessed clot stiffness, reflecting clots’ resistance to deformation, see Figure 2C. Clots from male subjects have significantly lower stiffness than clots from female subjects (4.78 ± 0.43 vs. 6.28 ± 0.91 kPa, p = 0.013). We then quantified fracture toughness, which describes a material’s resistance to crack propagation, using Equation 1, see Figure 2D. Clots from male subjects have a significantly lower fracture toughness than clots from female subjects (1.46 ± 0.08 vs. 2.19 ± 0.33 J/m^2^, p = 0.001). Next, we measured clot strength as the peak stress of pure shear samples, see Figure 2E. Clots from male and female subjects do not significantly differ in strength (2.82 ± 0.49 vs. 3.89 ± 0.94 kPa, p = 0.064). Finally, we calculated work to rupture, representing the total energy absorbed by the clot prior to failure. Work to rupture was not significantly different between clots from male and female subjects (0.55 ± 0.12 vs. 0.82 ± 0.23 kJ/m^3^, p = 0.064).

**Figure 2:**
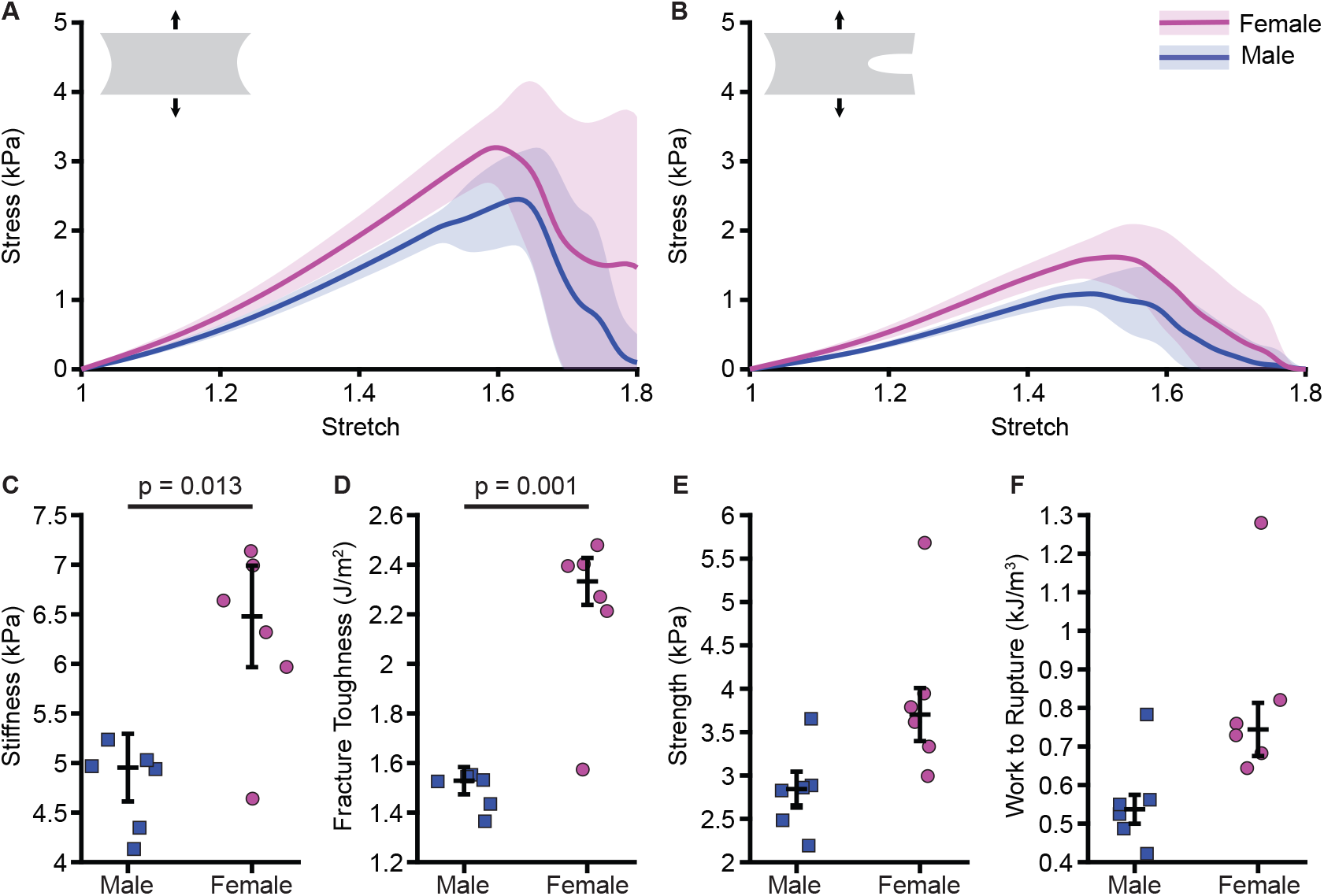
Blood clot mechanics by sex. (A-B) Stress-stretch curves differ between clots from male and female subjects in both pure shear and mode-I tests. Curves show mean ± 1 standard deviation. (C-D) Clots from male subjects have lower stiffness and fracture toughness than clots from female subjects. (E-F) There are no significant sex differences in clot strength and work to rupture. Data points represent the mean of three technical replicates per subject. Error bars show mean ± 1 standard deviation.

### 3.3. Blood clot mechanical properties are associated with multiple blood composition metrics

Figure 3 summarizes correlations between clot mechanical properties and blood composition metrics. We found that mechanical properties are most strongly associated with red blood cell–related metrics and with fibrinogen. Higher hematocrit, hemoglobin, and RBC counts are associated with decreased stiffness, fracture toughness, strength, and work to rupture. In contrast, higher fibrinogen concentrations correlate with increases in all four properties. Most white blood cell measures do not show meaningful relationships with clot mechanical properties. However, we observe some weaker associations, including positive correlations between neutrophils and strength and work to rupture. Conversely, eosinophils negatively correlate with strength. We do not detect significant relationships for total WBC count, platelets, lymphocytes, monocytes, or basophils.

**Figure 3:**
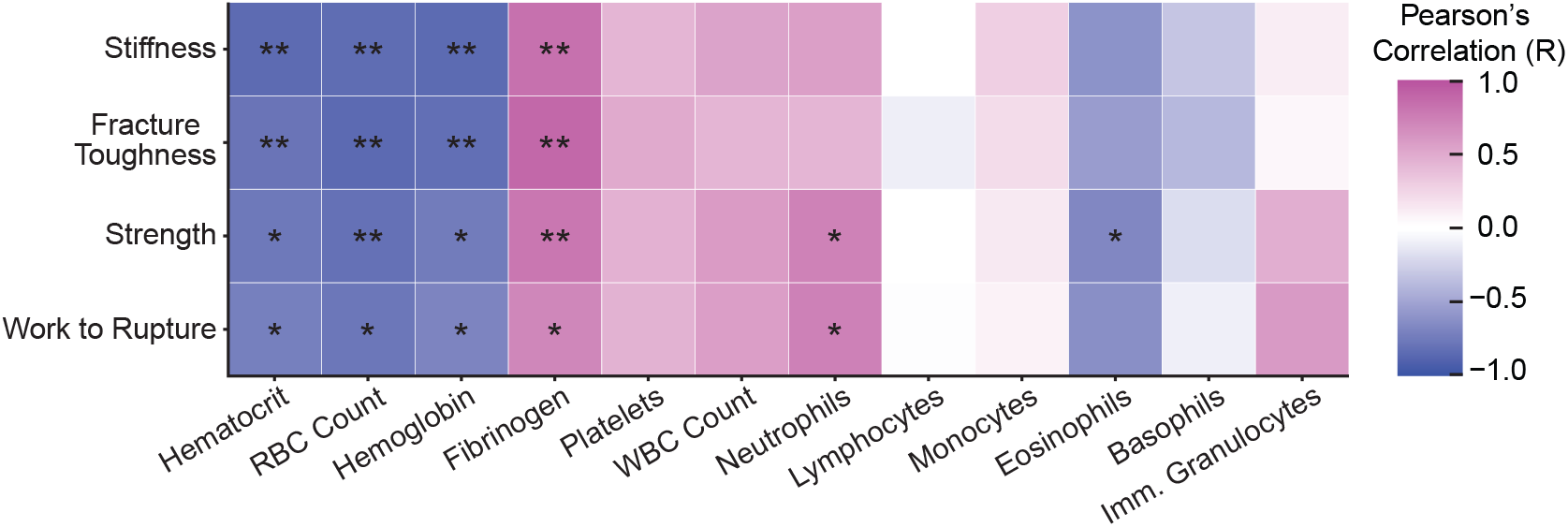
Heatmap showing Pearson’s correlations between blood clot mechanical properties (stiffness, fracture toughness, strength, work to rupture) and CBC measures. Color indicates the correlation coefficient (R), with blue representing negative correlations, pink representing positive correlations, and white near zero. Asterisks indicate statistical significance (*p < 0.05, **p < 0.01).

Figure 4 highlights the specific blood composition metrics that showed significant relationships with clot mechanical properties. These linear regressions, along with 95% confidence intervals, illustrate the direction and strength of the associations. Since hematocrit, hemoglobin, and RBC count are highly correlated measures of blood composition, we used hematocrit as a representative metric for this group. Hematocrit accounts for a large portion of the observed variation in stiffness (R^2^ = 0.72, p = 0.006), fracture toughness (R^2^ = 0.63, p = 0.01), strength (R^2^ = 0.59, p = 0.014), and work to rupture (R^2^ = 0.52, p = 0.023), see Figure 4A. Similarly, fibrinogen explains a substantial amount of variation in stiffness (R^2^ = 0.70, p = 0.006), fracture toughness (R^2^ = 0.78, p = 0.006), strength (R^2^ = 0.64, p = 0.01), and work to rupture (R^2^ = 0.50, p = 0.03), see Figure 4B. Neutrophils contribute moderately to the observed variation in strength (R^2^ = 0.54, p = 0.02) and work to rupture (R^2^ = 0.55, p = 0.019), whereas eosinophils had the smallest explanatory power, yielding a significant regression with strength (R^2^ = 0.45, p = 0.043), see Figure 4C and D.

**Figure 4:**
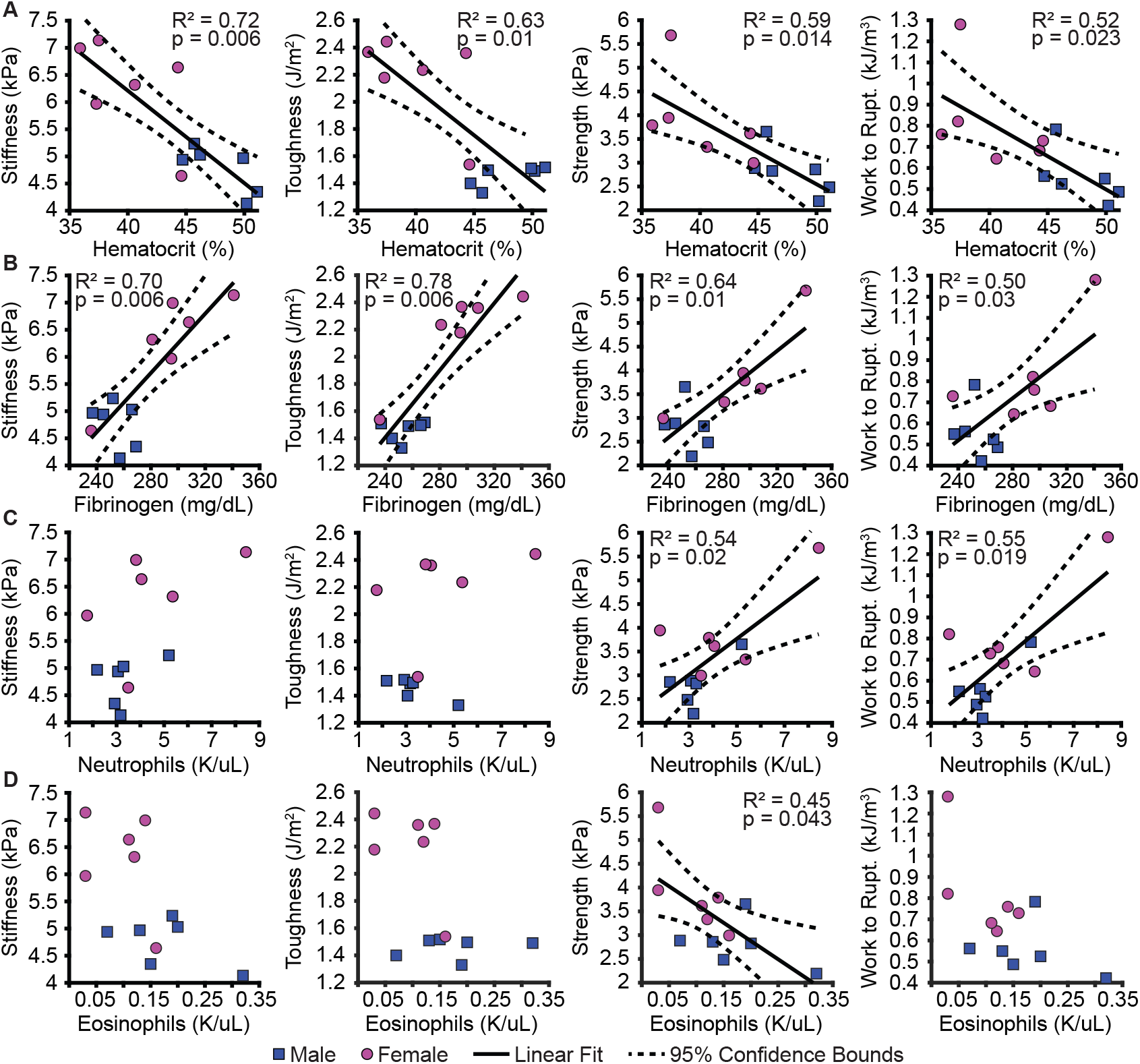
Linear regressions between clot mechanical properties and select CBC measures. (A) Hematocrit shows a negative relationship with all four mechanical metrics. (B) Fibrinogen is positively associated with all four metrics. (C) Neutrophil count positively correlates with strength and work to rupture. (D) Eosinophil count negatively correlates with strength. Data points represent the mean of three technical replicates per subject. Linear regression fits and 95% confidence intervals are indicated for statistically significant correlations.

## 4. Discussion

Our goal was to identify what blood components drive variability in the mechanical properties of in vitro blood clots. To this end, we drew blood from a young and healthy adult population, coagulated clot samples for mechanical testing, and measured subjects’ blood composition. We then tested how clot stiffness, fracture toughness, strength, and work to rupture depend on blood composition measures. In summary, we found that clot mechanical properties are most strongly associated with sex, hematocrit, fibrinogen, neutrophils, and eosinophils. We did not observe significant relationships with platelets and other WBC sub-types.

More specifically, we first found that blood composition and blood clot mechanical properties differ between male and female subjects. Blood from female subjects has lower hematocrit, RBC counts, and hemoglobin compared to male subjects. Furthermore, clots from female subjects are stiffer and have a higher fracture toughness than clots from male subjects. These findings are consistent with our prior work, which examined a larger and more heterogeneous subject population [10]. Practically, these data show that clots from female subjects are less deformable and more resistant to rupture than those from male subjects. These differences likely influence how clots embolize and obstruct vessels [1, 18, 19]. However, whether they account for sex differences in thromboembolic disease remains to be determined [20–23]. Further work is needed to directly link sex differences in clot mechanical properties to clinical outcomes.

Second, we found that clot mechanical properties correlate strongly with RBC measures, including hematocrit, RBC count, and hemoglobin. Specifically, as hematocrit increases, clot mechanical properties decrease. Interestingly, this finding differs from our prior study, in which stiffness showed no relationship with hematocrit and fracture toughness exhibited only a weak negative correlation [10]. The stronger association observed in the present study likely reflects our narrow age range, which minimizes confounding effects of age-related changes in blood composition, hormone levels, and metabolic variability [24–28]. In fact, our results align with studies that directly manipulate hematocrit across a wide range to assess its impact on clot mechanics, confirming that hematocrit is a key determinant [5, 29].

Third, we found that fibrinogen shows strong positive correlations with clot mechanical properties. It is important to note that fibrinogen and hematocrit are inversely correlated in our data (R = -0.67, p = 0.016), and both have high explanatory power. However, fibrinogen accounts for the largest proportion of variability, on average, across all four metrics. Fibrinogen explains 70% of stiffness, 78% of fracture toughness, 64% of strength, and 50% of work to rupture. To our knowledge, no prior study has directly examined the impact of physiological blood fibrinogen levels on whole blood clot mechanics. However, our findings align with in vitro studies on fibrin gels and plasma clots, where increased fibrinogen concentration produces denser, stronger fibrin networks [30–32]. Clinically, fibrinogen is known to be critical for hemostasis, and supplementation is used to improve clot strength and stability in trauma or surgical patients [33–36]. Our findings show that, even among healthy individuals, physiological variations in fibrinogen are associated with differences in clot mechanical properties. Thus, fibrinogen is critical to clot mechanics across both pathological and normal physiological conditions.

Finally, we found that some WBC subtypes are associated with clot mechanical properties. Our prior work suggested that total WBC count affected blood clot mechanical properties. However, without differential WBC counts, the contributions of individual subtypes were unclear. Here, we identify neutrophils and eosinophils as key contributors. Interestingly, they have opposing effects: neutrophils positively correlate with clot strength and work to rupture, whereas eosinophils negatively correlate with strength. This finding aligns with prior in vitro studies showing that neutrophils interact with fibrin networks and enhance clot stability [37]. They do so by forming neutrophil extracellular traps (NETs), which structurally reinforce the fibrin network [38, 39]. In contrast, the role of eosinophils in clot mechanics is less well-characterized. In vivo, eosinophils release toxic granule proteins that can disrupt extracellular matrix architecture [40, 41]. A similar mechanism may occur in clots, where eosinophils interfere with the fibrin network structure and weaken clots. However, future work is needed to test the mechanisms by which eosinophils affect fibrin network structure and clot mechanical properties.

Our study has limitations. First, our in vitro clots are a simplified model of in vivo clots. They are homogeneous, rich in RBCs, formed without flow, and lack the collagen deposits of aged thrombi [42, 43]. This makes our model most representative of fresh venous clots. Future studies should investigate a broader range of clot compositions. Second, our study population was small and relatively homogeneous. While this yielded strong relationships between mechanical properties and blood composition measures, it also raises the question of whether these findings hold across more diverse populations. Finally, we did not account for sex-specific factors beyond biological sex, such as hormonal cycles, which could influence clot properties [44–46]. Future studies should extend these findings by including larger, more heterogeneous populations and exploring additional physiological variables.

In summary, we found that clot mechanical properties are strongly influenced by sex, RBC measures, fibrinogen levels, and specific WBC subtypes, including neutrophils and eosinophils. To our knowledge, this is the first study to demonstrate that physiological variations in fibrinogen and individual WBC subtypes affect whole blood clot mechanics. These findings expand our understanding of the biological factors that contribute to clot heterogeneity and provide new insight into how blood composition may influence thromboembolic risk and treatment response. Future work should aim to directly link in vitro clot mechanical properties with clinical outcomes in thromboembolic disease to better understand their physiological and pathological relevance.

## Author Contributions

G.N.B. designed and performed experiments, analyzed data, and drafted the manuscript. N.S. performed experiments. J.N. and T.E. analyzed data. J.F., J.T., H.S., and S.H.P. contributed to study conceptualization and data interpretation. M.K.R. contributed to study conceptualization and design, data interpretation, and manuscript writing. All authors revised and approved the final manuscript.

## Relationship Disclosure

Manuel K. Rausch has a speaking agreement with Edwards Lifesciences. The other authors have no conflicts to declare.

## Data Availability

Data are available upon request.

