## Supplemental Information for "In vitro blood clot mechanical properties depend on fibrinogen and white blood cell subtypes in addition to hematocrit"

**Table of Contents:**

- Table S1

| Subject | Sex | Age | WBC | RBC | Hemoglobin | Hematocrit | MCV | MCH | MCHC | Platelets | RDW |
| --- | --- | --- | --- | --- | --- | --- | --- | --- | --- | --- | --- |
| 1 | F | 22 | 9.7 | 4.78 | 13.2 | 40.6 | 84.9 | 27.6 | 32.5 | 340 | 11.9 |
| 2 | F | 24 | 7.4 | 4.56 | 14.8 | 44.3 | 97.1 | 32.5 | 33.4 | 271 | 12.2 |
| 3 | F | 23 | 6.4 | 4.19 | 11.7 | 35.9 | 85.7 | 27.9 | 32.6 | 262 | 13.1 |
| 4 | F | 25 | 5.7 | 4.76 | 15 | 44.6 | 93.7 | 31.5 | 33.6 | 251 | 12 |
| 5 | F | 25 | 3.2 | 3.95 | 12.2 | 37.3 | 94.4 | 30.9 | 32.7 | 245 | 11.9 |
| 6 | F | 25 | 10.8 | 4.02 | 12.2 | 37.5 | 93.3 | 30.3 | 32.5 | 324 | 11.7 |
| 7 | M | 26 | 5.6 | 5.46 | 16.9 | 50.2 | 91.9 | 31 | 33.7 | 233 | 12.3 |
| 8 | M | 27 | 5.6 | 5.52 | 16.8 | 51.1 | 92.6 | 30.4 | 32.9 | 316 | 11.9 |
| 9 | M | 25 | 5.1 | 5.16 | 14.7 | 44.7 | 86.6 | 28.5 | 32.9 | 227 | 12.4 |
| 10 | M | 24 | 5.8 | 5.23 | 15.5 | 46.2 | 88.3 | 29.6 | 33.5 | 202 | 12.4 |
| 11 | M | 22 | 8.2 | 5.21 | 16 | 45.7 | 87.7 | 30.7 | 35 | 276 | 13.7 |
| 12 | M | 20 | 4.6 | 5.35 | 16.2 | 49.9 | 93.3 | 30.3 | 32.5 | 201 | 11.8 |

| **Subject** | **Neutrophils** | **Lymphocytes** | **Monocytes** | **Eosinophils** | **Basophils** | **IG** | **NRBCs** | **Fibrinogen** |
| --- | --- | --- | --- | --- | --- | --- | --- | --- |
| 1 | 5.36 | 0.5 | 0.62 | 0.12 | 0.06 | 0.02 | 0 | 281 |
| 2 | 4.05 | 2.72 | 0.49 | 0.11 | 0.04 | 0.02 | 0 | 308 |
| 3 | 3.82 | 1.87 | 0.55 | 0.14 | 0.02 | 0 | 0 | 296 |
| 4 | 3.49 | 1.57 | 0.36 | 0.16 | 0.06 | 0.03 | 0 | 236 |
| 5 | 1.76 | 1.18 | 0.22 | 0.03 | 0.01 | 0.01 | 0 | 295 |
| 6 | 8.44 | 1.69 | 0.52 | 0.03 | 0.04 | 0.04 | 0 | 341 |
| 7 | 3.17 | 1.53 | 0.54 | 0.32 | 0.06 | 0.02 | 0 | 257 |
| 8 | 2.92 | 2.03 | 0.42 | 0.15 | 0.03 | 0.01 | 0 | 269 |
| 9 | 3.06 | 1.53 | 0.38 | 0.07 | 0.04 | 0.01 | 0 | 245 |
| 10 | 3.29 | 1.73 | 0.51 | 0.2 | 0.03 | 0.01 | 0 | 266 |
| 11 | 5.2 | 2.09 | 0.57 | 0.19 | 0.07 | 0.03 | 0 | 252 |
| 12 | 2.18 | 1.97 | 0.23 | 0.13 | 0.04 | 0.01 | 0 | 237 |

**Table S1** Summary of subjects’ sex, age, and blood composition data. Complete blood count data includes white blood cell (WBC) count (K/uL), red blood cell (RBC) count (M/uL), hemoglobin (g/dL), hematocrit (%), mean cell volume (MCV, fL), mean corpuscular hemoglobin (MCH, pg), mean corpuscular hemoglobin concentration (MCHC, g/dL), platelet count (K/uL), red cell distribution width (RDW, %), neutrophil count (K/uL), lymphocyte count (K/uL), monocyte count (K/uL), eosinophil count (K/uL), basophil count (K/uL), immature granulocytes (IG, K/uL), nucleated RBCs (NRBCs, K/uL), and fibrinogen levels (mg/dL).
